# Bioelectromagnetic platform for stimulation

**DOI:** 10.1101/2021.06.07.447412

**Authors:** Ryan C Ashbaugh, Lalita Udpa, Ron R. Israeli, Assaf A. Gilad, Galit Pelled

## Abstract

Magnetogenetics is a new field that utilizes electromagnetic fields to remotely control cellular activity. In addition to the development of the biological genetic tools, this approach requires designing hardware with a specific set of demands for the electromagnets used to provide the desired stimulation for electrophysiology and imaging experiments. Here we present a universal stimulus delivery system comprised of four magnet designs compatible with electrophysiology, fluorescence and luminescence imaging, microscopy, and freely behaving animal experiments. The overall system includes a low-cost stimulation controller which enables rapid switching between active and sham stimulation trials as well as precise control of stimulation delivery.

## 1. Introduction

The rapid growth of research interest in magnetogenetics in the past decade has resulted in a broad range of bio-electromagnetic stimulation applications (Nimpf and Keays, 2017), creating a demand for sophisticated stimulus delivery systems. Many biological systems can be magnetically stimulated to regulate gene expression or neural activity, and stimulation parameters can vary significantly depending on the mechanisms employed to elicit responses (Nimpf and Keays, 2017). In contrast to visible light, low frequency and DC magnetic fields easily penetrate soft tissue and bone, potentially allowing for minimally invasive and wireless stimulation. High costs of bioelectromagnetic stimulation devices and a lack of systematic analysis of electromagnetic stimulus fields serve as a barrier to designing quantitative studies and replicating results in magnetogenetics experiments.

Development of magnetic sensitive pathways, like those using nanoparticles (Chen et al., 2015) and proteins like the electromagnetic perceptive gene (EPG) (Krishnan et al., 2018; Mitra et al., 2020; Cywiak et al., 2020; Hwang et al., 2020; Hunt et al., 2021) have contributed to making magnetic stimulus delivery for wide ranging applications increasingly important. Furthermore, recent studies which show that humans may also have magnetoperception (Wang et al., 2019) serve to increase the demand for easy to implement and versatile electromagnetic stimulation devices.

One solution for magnetic stimulus delivery includes the use of an induction heater (Chen et al., 2015), however these are limited in their ability to be integrated into a wide variety of experimental protocols. Therefore, it is beneficial to design and build electromagnets which can be more easily incorporated into a wide variety of applications. Custom stimulation coils demonstrate improved integration into microscopy applications (Pashut et al., 2014; Hernández-Morales et al., 2020), and we aim to build on this flexibility and emphasize detailed stimulation validation. Further, repeatability, uniformity, a negative control condition, and ease of use are critical properties of interest in magnetic stimulus systems.

Here we present work conducted toward developing a magnetogenetics bioelectromagnet stimulation platform which is low cost, versatile, easy to use and affords a high degree of control over stimulation parameters. The electromagnet designs presented in this paper are applicable for electrophysiology, microscopy, fluorescence and luminescence imaging, and also to stimulate in freely behaving animals.

We developed four designs, where each design is unique to accommodate application specific physical constraints as well as maintain uniformity in the target area. Double wrapping coils as described in Kirschvink et al. (Kirschvink, 1992) allows experiments to be tested with a negative control and the use of a low-cost stimulation controller. An additional graphical interface provides a user-friendly way to switch between active and sham stimulation conditions and reproduce specific stimulus parameters.

## 2. Applications

The primary use of the electromagnet systems we designed are microscopy, luminescence and fluorescence imaging, *in vivo* electrophysiology and freely behaving animal experiments. For each application our goal was to design an electromagnet that can deliver the desired magnetic flux given various constraints including power consumption, coil and sample temperature, and coil size. Evidence suggests that applying magnetic flux densities >50 mT (Wheeler et al., 2016; Krishnan et al., 2018; Hunt et al., 2021) was successful at eliciting responses in magnetoreceptive targets. Thus, this paper presents a system that provides the ability to conduct a parametric study of potential stimulation parameters and investigate the response thresholds of these parameters.

### 2.1. Microscopy

Microscopy applications tend to impose strict size constraints on electromagnets. As also seen in Pashut et al. (Pashut et al., 2014), care must be taken to ensure that the electromagnet does not interfere with the objectives or condenser of a microscope. For fluorescence microscopy, calcium imaging, voltage imaging, and patch clamping, a single coil was designed specifically to fit around a circular 35 mm diameter glass bottomed cell culture dish. During imaging, the electromagnet is either placed within a microscope compatible auxiliary incubation chamber or mounted underneath the stage directly below the sample, depending on the microscope in use.

The incubation chamber restricts the maximum width and height of the coil holder to 85 mm and 15 mm respectively. An assembled coil placed around a 35 mm culture dish is illustrated schematically in Figure 1a, where only half of the coil is shown for clarity. An advantage of this design, as will be shown in Section 3, is that the field in the target region at the center of the plate is relatively uniform thereby providing the ability to reliably deliver consistent stimulus between experiments.

**Figure 1:**
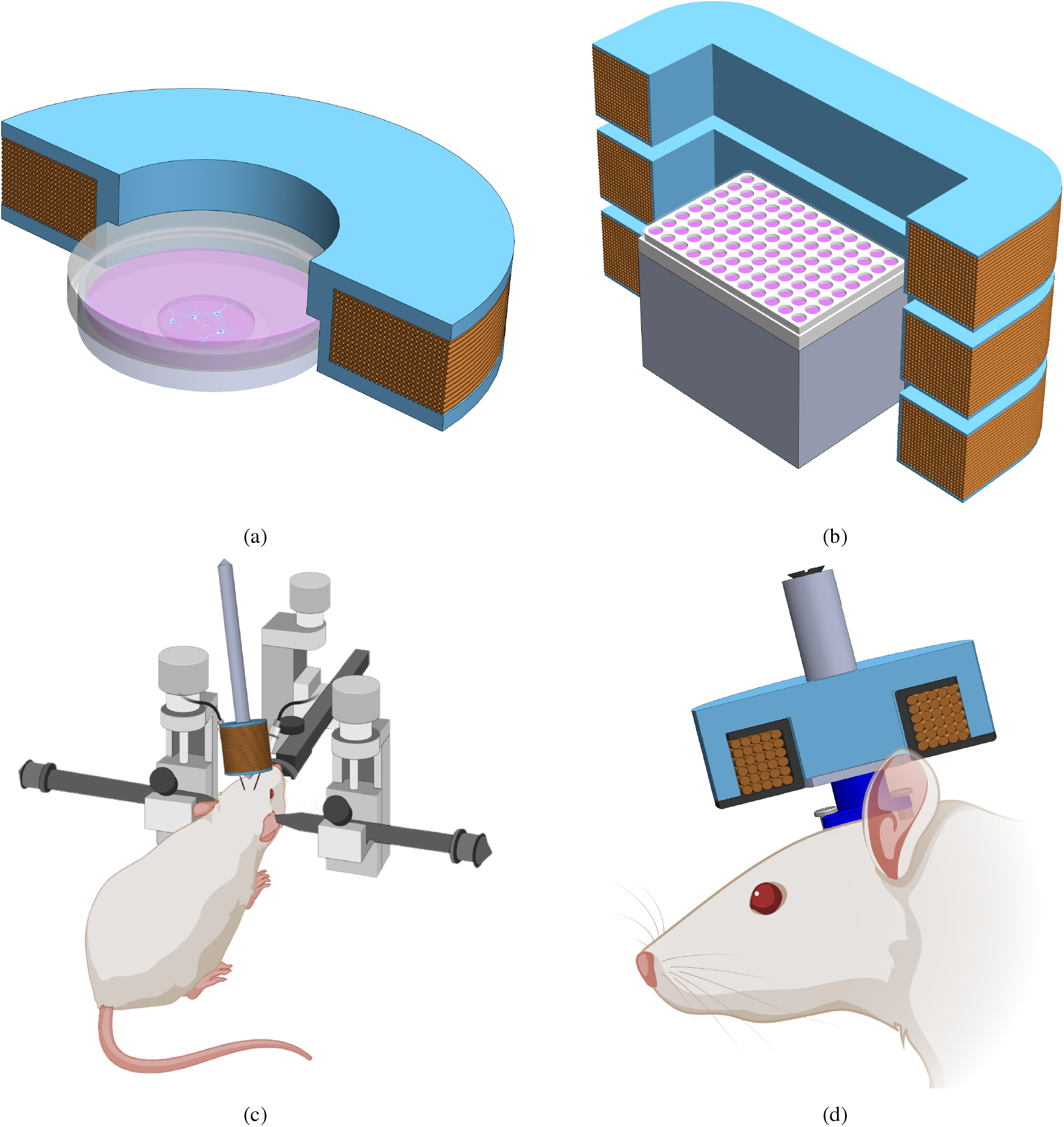
Stimulation applications are pictured for each of the four geometries under study, where (a), (b) and (d) show half of a coil for improved visualization. (a) shows a coil designed to fit overtop of a 35 mm culture dish which can be used for microscopy. (b) shows a set of three-coils which can be stacked and used for stimulating samples on a multiwell plate with a relatively uniform field. (c) demonstrates an application of an electromagnet for head-fixed electrophysiology. (d) pictures a coil wound onto a bobbin and inserted into a pot core which could be used for freely behaving animal studies. (Biorender, 2021)

### 2.2. Luminescence and Fluorescence Imaging

This application consists of measuring responses to magnetic stimuli in many cell preparations at the same time. A multi-well plate is used to test the effects of cell type, media preparation, control conditions, or genetic variants of a protein all within the same trial. This enables higher throughput for screening experiments, opening the door for mutagenesis studies aimed at improving stimulation responses. The application requires consistent stimulus delivery in each trial. To facilitate higher throughput screening, a three-coil electromagnet was designed based on the Merritt Coils outlined in (Merritt et al., 1983; Kirschvink, 1992; Magdaleno-Adame et al., 2010) and used for stimulation of multi-well plates within a PerkinElmer In Vivo Imaging System (IVIS).

Using a multi coil design aids in producing a uniform magnetic field within a volume along the central axis of the coil. Figure 1b shows an illustration of half of the three-coils with a 96 well plate placed at the central plane.

### 2.3. In Vivo Electrophysiology

The potential neuromodulation effects of magnetic stimulation on rodents expressing the EPG protein or any other magnetoreceptive gene is considered in this application. An electromagnet design consisting of a coil wound around a ferromagnetic core is proposed. This coil was attached to an adjustable arm and positioned next to the head of a rat while recording neural signals.

This design is particularly useful when the application allows for less restrictions on the physical placement of the coil. A schematic of the experimental setup for this application is presented in Figure 1c, where the electromagnet is positioned between electrodes placed in the brain of an anesthetized, head-fixed rat. Typically, there is much more space to place the equipment and adjust it for proper alignment with the target in electrophysiology than in applications such as microscopy, making this a versatile solution.

### 2.4. Freely Moving Animal

In addition to the aforementioned methods of investigating magnetosensitive pathways, it is also of great benefit to be able to study the effects of stimulation on the behavior of freely moving animals. Such studies could be performed in an operant conditioning box and designed to monitor reward seeking behavior, anxiety, stress, etc.

This application, however, presents a challenge for stimulus delivery. In the case of rodents, cages used for behavioral studies can vary from 200-500 mm in length and width, and could be as tall as 300 mm. While Merritt Coils can deliver uniform stimulus to a given volume, for delivering uniform fields to a large volume, the required power can exceed the capabilities of practical systems. Delivering stimulus of up to ~50 mT within a uniform field in a region the size of a rodent cage using Merritt Coils is therefore impractical.

Alternatively, a stimulus can be delivered locally using a fixed stimulation device attached to the subject. The attachment may consist of a head mounted fixture or a wearable jacket and would allow the animal to freely move about. Stimulus can then be applied to the localized area when desired. For this application, an electromagnet built into a pot core provides a good solution. Pot cores consist of a central rod around which a coil is wrapped, and a hi-mu metal shield surrounding the coil. The hi-mu metal is highly permeable to magnetic fields and serves to increase magnetic flux density and focus the magnetic fields within the core region. A depiction of such a device is shown in Figure 1d, where it is attached to a custom head mounting fixture.

## 3. Numerical Modeling

Before fabricating and assembling the electromagnets and associated components, each design was numerically modeled and simulated using finite element analysis to assess the performance of the design. Simulations were implemented using COMSOL Multiphysics (Multiphysics, 2014) to solve the electromagnetic field equations governing the underlying physics. The visualization of the magnetic flux distribution allows for optimization of the geometric parameters of the design. Unless otherwise specified, all magnetic flux density simulations were performed with a constant volumetric current of 15 Amperes passing through the coil.

### 3.1. Air Core Model

A single coil geometry was modeled to fully utilize the space available within the imaging incubation chamber for a Keyence BZ-X800E microscope. A coil with 264 turns and height of 11.5 mm, outer diameter of 85.0 mm, and inner diameter of 45.0 mm was considered. Figure 2a shows a schematic of the simulated coil and Figure 2b shows the simulation results of the magnetic flux density magnitude in the YZ plane. Figure 2c shows the simulated line scans along the lines depicted in Figure 2b, showing that stimulations greater than 50 mT are expected.

**Figure 2:**
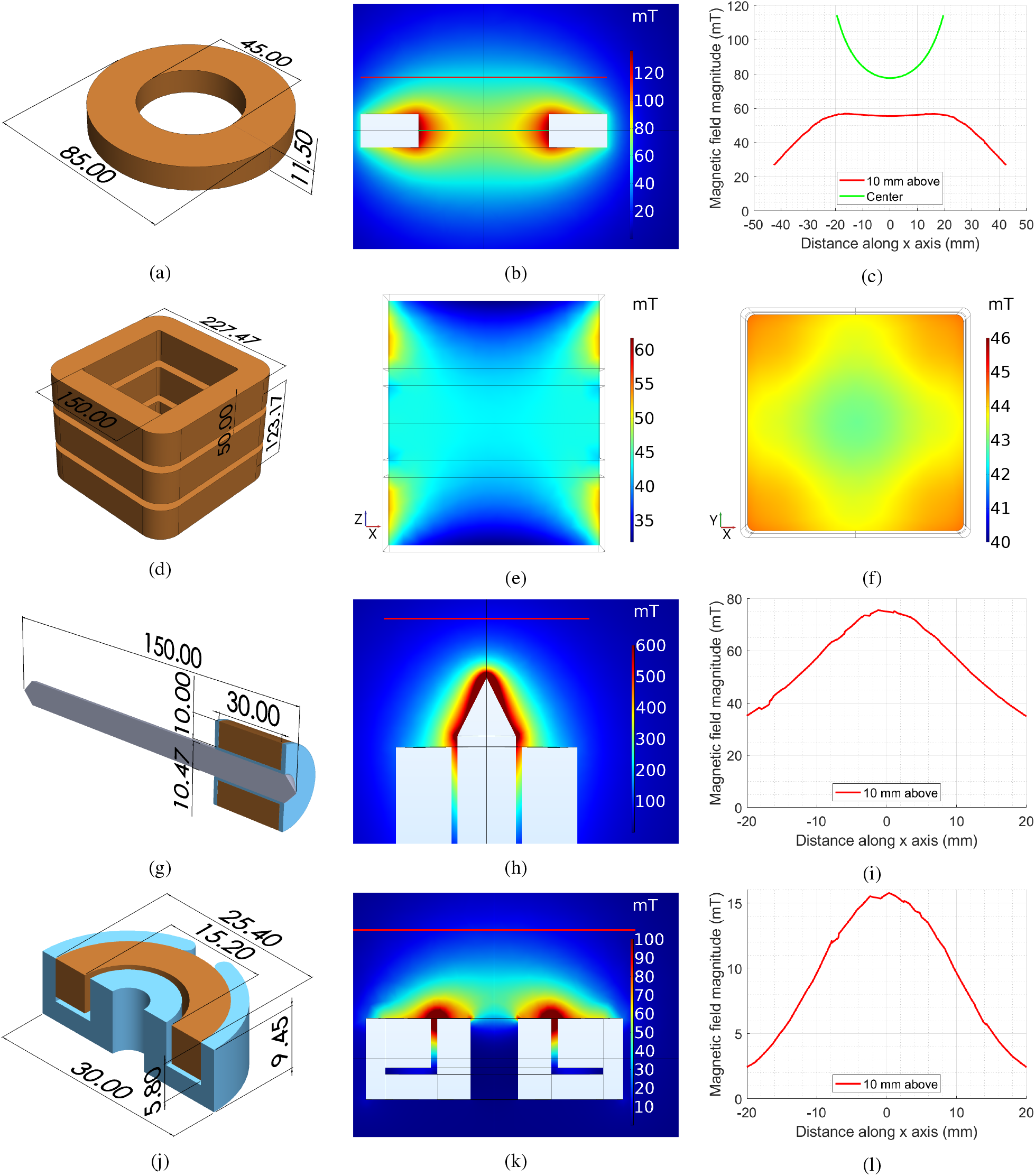
Electromagnet geometries and simulated magnetic flux density magnitude. Dimensions for the (a) air core coil, (d) three-coil system, (g) ferromagnetic core coil and (j) pot core coil. Simulated magnetic flux density magnitude for the (b) air core coil, (e) three-coil system XZ plane, (f) three-coil system XY plane, (h) ferromagnetic core coil, and (k) pot core coil. The superimposed lines represent the location of the line scans (c) for the air core coil, (i) ferromagnetic core coil, and (l) pot core coil.

### 3.2. Three-Coil System Model

A three-coil geometry was modeled to fit inside of an IVIS having an imaging chamber of dimensions 430×380×430 mm. Due to a 100 V supply voltage constraint and 15 Ampere supply current constraint, the total device resistance was constrained to 6.66 n. The coil was modeled with 14 gauge magnet wire, selected due to its low resistivity. The maximum length of the wire was determined based on the maximum resistance and the resistivity.

Figure 2d shows a representation of the dimensions of the three-coil system. 150 mm was selected for the inner square side length since the multi-well plates are 130 mm wide. The total height of the system was chosen to be 123.17 mm, consistent with the ratio of side length to coil spacing presented in (Merritt et al., 1983) for a three-coil Merritt Coil system,

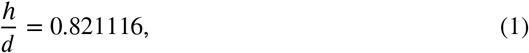

where d is the length of each side of the coils and h is the height. The height and outer width of each coil was selected to be 50 mm and 227.47 mm respectively, resulting in a simulated coil thickness of 38.75 mm and 276 turns per coil.

A side view of the central XZ plane is shown in Figure 2e having a magnetic flux density magnitude of about 45 mT in the center. Figure 2f shows the central XY plane, which clearly demonstrates the uniformity of the field in a central circular region of about 80 mm in diameter.

In contrast to coils shown in (Merritt et al., 1983; Kirschvink, 1992; Magdaleno-Adame et al., 2010) which use an ampere turn ratio of 0.512 for the center coil relative to the top and bottom coils, the system modeled utilizes coils of equal ampere turn ratios which adds more turns to the system. A central coil with ampere turn ratio 20:39 can be substituted should a volume of uniformity be required instead of a plane.

### 3.3. Ferromagnetic Core Model

A ferromagnetic core of diameter 10.47 mm, length 150 mm, and relative magnetic permeability of 6,500 was modeled. The tips at each end are tapered to a point as seen in Figure 2g. The coil geometry with an outer diameter of 31.97 mm wrapped along 30 mm of the length of the core. The coil was simulated with 372 turns.

Figure 2h shows that the field exceeds 600 mT at the very tip of the core. This flux then decays quickly along the coil axis, dropping to about 75 mT at a distance of 10 mm from the tip of the core, as seen in Figure 2i.

### 3.4. Pot Core Model

Lastly the pot core configuration for freely moving animals was simulated. For a pot core electromagnet to be head mounted on a rat, it must be small and light enough to allow maneuverability. A pot core geometry with an outer diameter of 30 mm and height of 9.45 mm was modeled along with the coil. A 28 turn coil was modeled to fit in the coil channel having depth and width of 6.5 mm and 6.05 mm respectively. The core was simulated with a high mu metal having a relative magnetic permeability of 10,000. Figure 2j shows the simulated pot core.

Since the device will be mounted on the moving animal, which is the target of stimulation, the magnetic field of interest is along the coil axis on the unshielded side of the pot core. The field distribution predicted by the numerical model is displayed in Figure 2k. While Figure 2k shows that there are strong fringing fields in close proximity to the coil, these fields decay quickly to yield uniform fields a few mm away. In practice, the coil holder and mounting device will cause the source to target distance to be a few mm. Figure 2l shows the magnetic field strength of the pot core along the coil axis at 10 mm from the device, indicating that a strength of ~15.5 mT at the target is achievable.

## 4. Magnet Assembly Implementation

### 4.1. Double Wrapping Coils

All coils pictured in Figure 3 are double wrapped, as demonstrated in (Kirschvink, 1992; Wang et al., 2019), to allow for a negative control. Wrapping a coil with two adjacent wires allows a user to reverse the direction of current in one wire relative to the other. This has the effect of cancelling out the magnetic fields generated by the two opposing currents, thereby resulting in a net zero field. The stimulus can then be operated in either active or sham mode, which can help to provide a control for motion caused by the changing magnetic flux or temperature increase due to ohmic heating in the coils.

**Figure 3:**
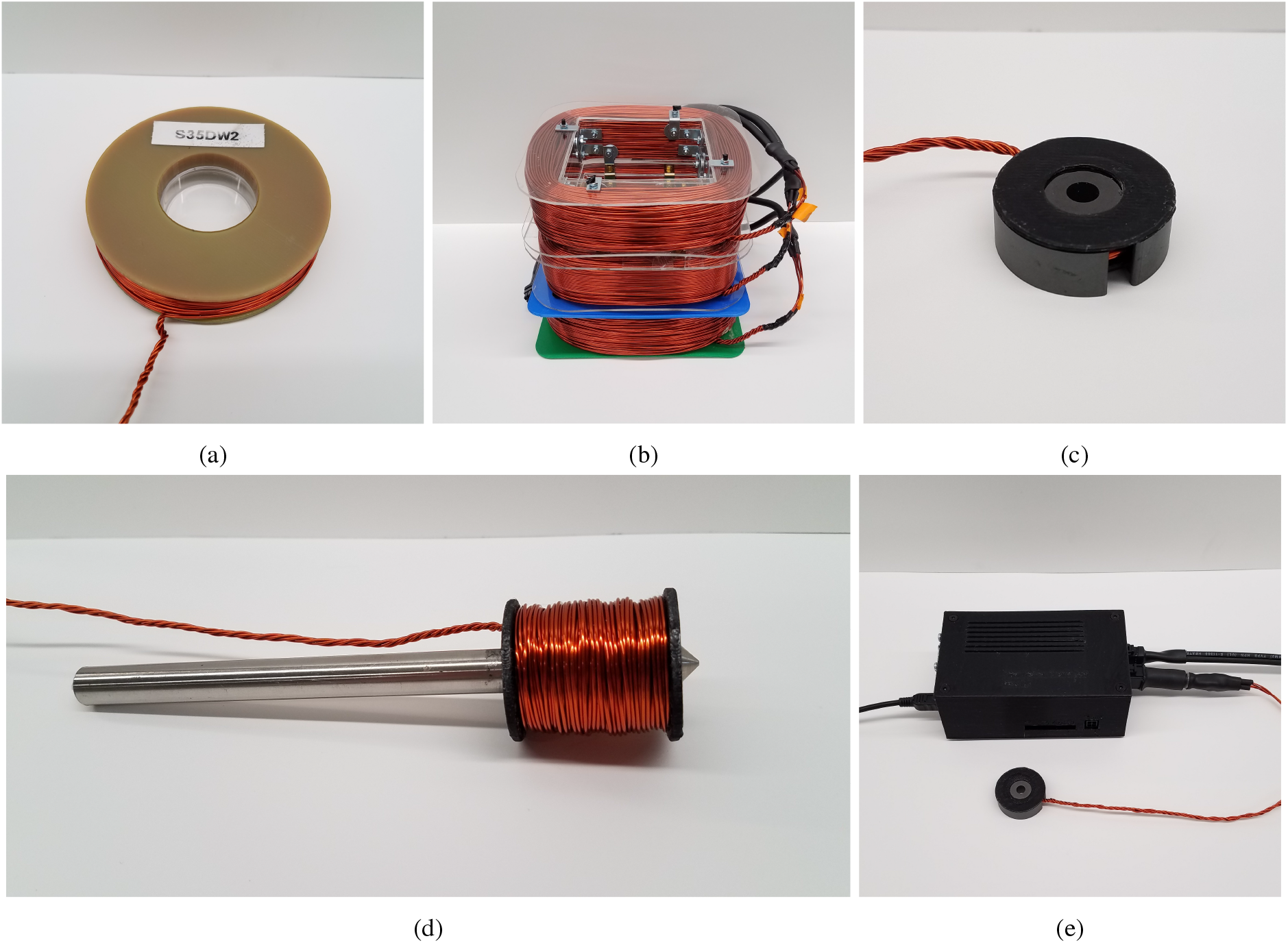
Assembled electromagnet devices. The air core coil in (a) is shown over top a 35 mm culture dish. (b) Shows the stacked three-coil system, where each coil is connected in series. (c) Pot core design shown with the unsheilded, open side up. (d) Ferromagnetic core coil. (e) The stimulation controller is pictured connected to a laptop with USB, the power supply which delivers the DC stimulation current, and the pot core coil.

In practice, it is not possible to fully cancel out the magnetic field in the sham mode because this would require perfectly aligned wires with negligible width. Regardless, empirical measurements of the magnetic field strength of the sham conditions discussed herein are typically at least an order of magnitude smaller than the field strength delivered in the active mode.

### 4.2. Air Core

A coil holder was 3D printed from high temperature plastic, heat treated, and then wrapped with 20 gauge wire such that the channel was evenly filled, resulting in 280 total turns. Figure 3a shows the assembled single coil placed over top of a 35 mm culture dish for comparison.

### 4.3. Three-Coil System

Three square coils were constructed to the same specifications used in the simulation model. The system is shown in Figure 3b with all three-coils together. Several methods were used in coil construction, with the initial method based on 3D printed parts. Due to the weight and size of the coils, the 3D printed parts were suboptimal in terms of strength and rigidity. Subsequent coil holders were assembled from acrylic components with metal hardware. While the top coil visible in Figure 3b is assembled with steel hardware, the central coil uses non-ferromagnetic brass hardware as described in (Wang et al., 2019).

### 4.4. Ferromagnetic Core

Two half bobbins were used when wrapping the coil around the ferromagnetic core so that the tightly wound wires would apply pressure on the half bobbins to maintain their position on the core. The total outer diameter of the coil is 30.7 mm, with an inner coil diameter equal to 10.7 mm, accounting for the core diameter and 3D printed bobbin. 308 turns of 20 gauge wire were used in the construction. Figure 3d shows the fully assembled ferromagnetic core electromagnet.

### 4.5. Pot Core

To get the maximum number of windings into the pot core channel, the coil is tightly wound around a 3D printed bobbin. Once wrapped, the bobbin is inserted into the pot core. A total of 28 turns of 20 gauge wire were fit into the channel. Figure 3c shows the assembled pot core device with the plastic bobbin inserted.

Attaching the pot core to the freely moving animal is also important for this application. To achieve secure attachment of the device while minimizing the distance between the coil and the stimulation target, a custom coil holder was designed to mount the pot core using the hole along the coil axis. This mount is itself attached to a permanent head mounted fixture that can be on the head of the animal. Alternatively, the device could also be attached to a wearable jacket to facilitate stimulation in other regions of an animal.

### 4.6. Stimulation Controller

One of the goals of this work was to make the electro-magnetic stimulation platform versatile and easy to use. Toward this goal, a stimulation controller was developed which allows the user to specify the stimulation protocol and switch between the active and sham conditions. Using automated stimulation protocols is highly advantageous as it increases the stimulation consistency between experiments.

A python application was developed to set the stimulation protocol and allow the user to control the stimulation delivered by a custom hardware device shown in Figure 3e, enabling selection of the direction of current through the double wrapped coils from within the graphical interface. Additionally, stimulations can be triggered by an external signal and an auxiliary stimulation signal can be connected to other devices.

## 5. Results and Discussion

It is crucial to validate simulated results with experimental measurements to fully characterize the core and coil parameters in an electromagnet stimulation system. For a quantitative comparison between simulation and experiment, we have utilized a 3-axis Gaussmeter along with a programmable XYZ scanner to measure the distribution of magnetic flux density in the regions of interest as shown in Figure 2 produced by the different configurations of stimulation devices.

All experimental measurements were performed with a 1 Ampere excitation current. At each position, 5 sensor measurements were averaged to generate the resulting magnetic flux density magnitude. To compare the experimental measurements with corresponding simulation results, magnetic flux density images were first aligned based on the maximum of their cross-correlation.

A qualitative comparison of the experimental and simulated field data is shown in Figure 4 for the air core, three-coil, ferromagnetic core, and pot core systems. The images represent spatial distribution of the magnitude of magnetic flux density. Results for the air core coil, shown in Figure 4a, indicate that a maximum magnetic flux density of 5.20 mT is recorded just above the surface of the coil. For the three-coil system, the first region of interest consists of the central XZ plane passing through the coils. A region of 60 mm by 120 mm centered at the center of the coil was scanned in the XZ plane, as shown in Figure 4b. In addition, measurements along the central XY plane were performed along a 40 mm by 120 mm region approximately centered on the coil axis, as shown in Figure 4c. In Figure 4d, we see the measured distribution of magnetic flux density for the ferromagnetic core coil, reaching a maximum of 24.55 mT. It is not surprising that of the four geometries, this design produces the highest magnetic flux density at the target location due to the effect of the ferromagnetic core. The pot core measurements are presented in Figure 4e, where the magnetic flux density achieves a maximum of 3.29 mT.

**Figure 4:**
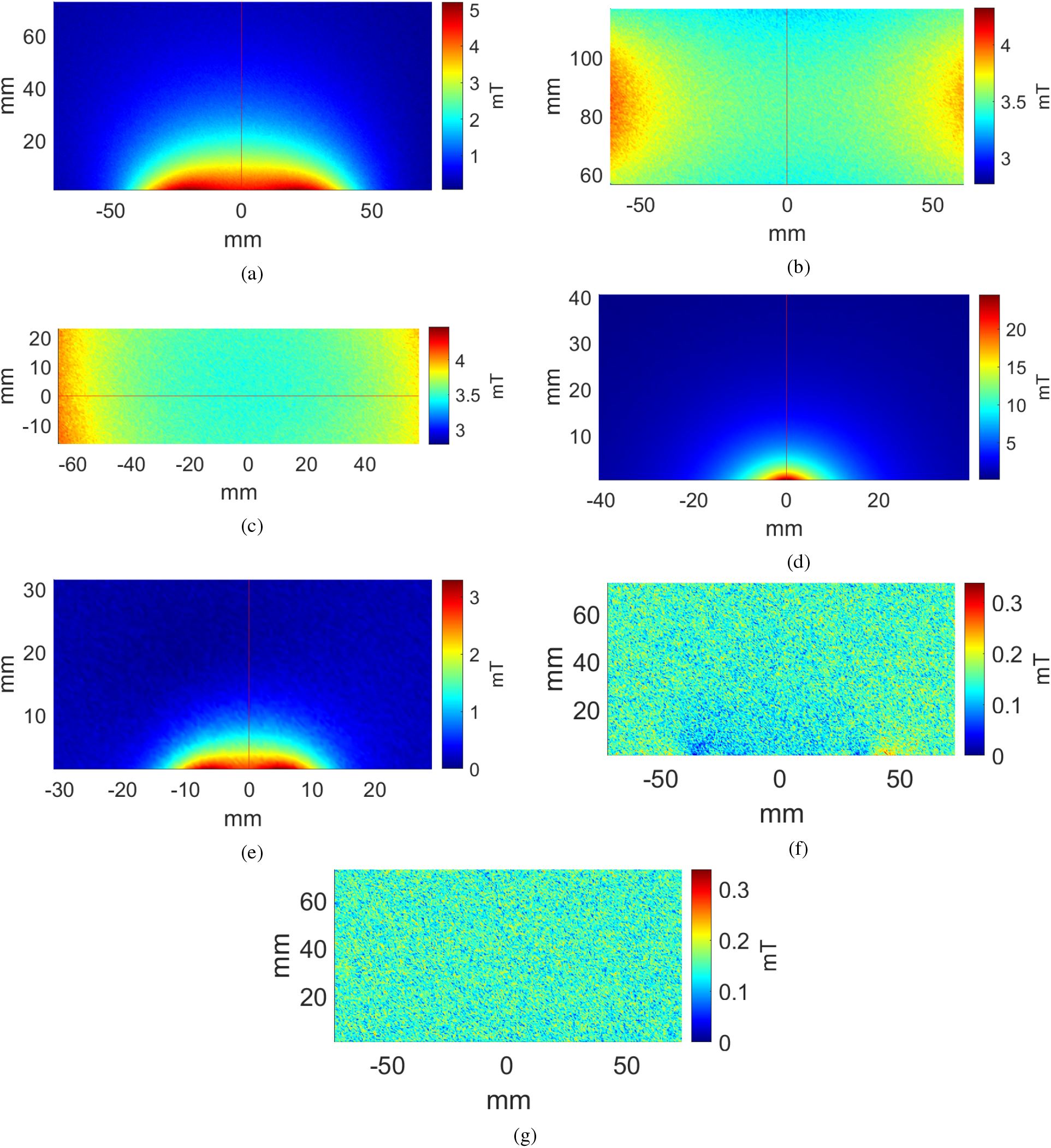
Experimental magnetic flux density magnitude distributions. (a) Air core coil active condition measurements along the central XZ plane. three-coil system active condition measurements along the central (b) XZ and (c) XY planes. (d) Ferromagnetic core coil active condition measurements along the central XZ plane. (e) Pot core coil active condition measurements along the central XZ plane. (f) Sham and (g) control condition measurements along the central XZ plane for the air core coil.

In addition to measuring the magnetic flux density magnitude during the active condition, similar measurements were taken in the sham configuration as well as with no stimulation current. The air core sham results are seen in Figure 4f, while the results for the air core no stimulation current control are shown in Figure 4g. Low amplitude fringing fields are observed near the coil at the bottom of the image in the case of the sham condition. Otherwise, the sham condition performed similarly to the no current case. Similar results were observed in the case of other geometries.

A quantitative comparison of the predicted and measured fields for the no current, sham, experimental and simulated stimulations for each coil design is shown in Figure 5 and summarized quantitatively in Figure 5f. The values listed in Figure 5f for the cases of air, ferromagnetic, and pot core coils are measured at a distance of 10 mm from the coil along the coil axis, while measurements for the three-coil system are taken at the center of the XZ and XY planes.

**Figure 5:**
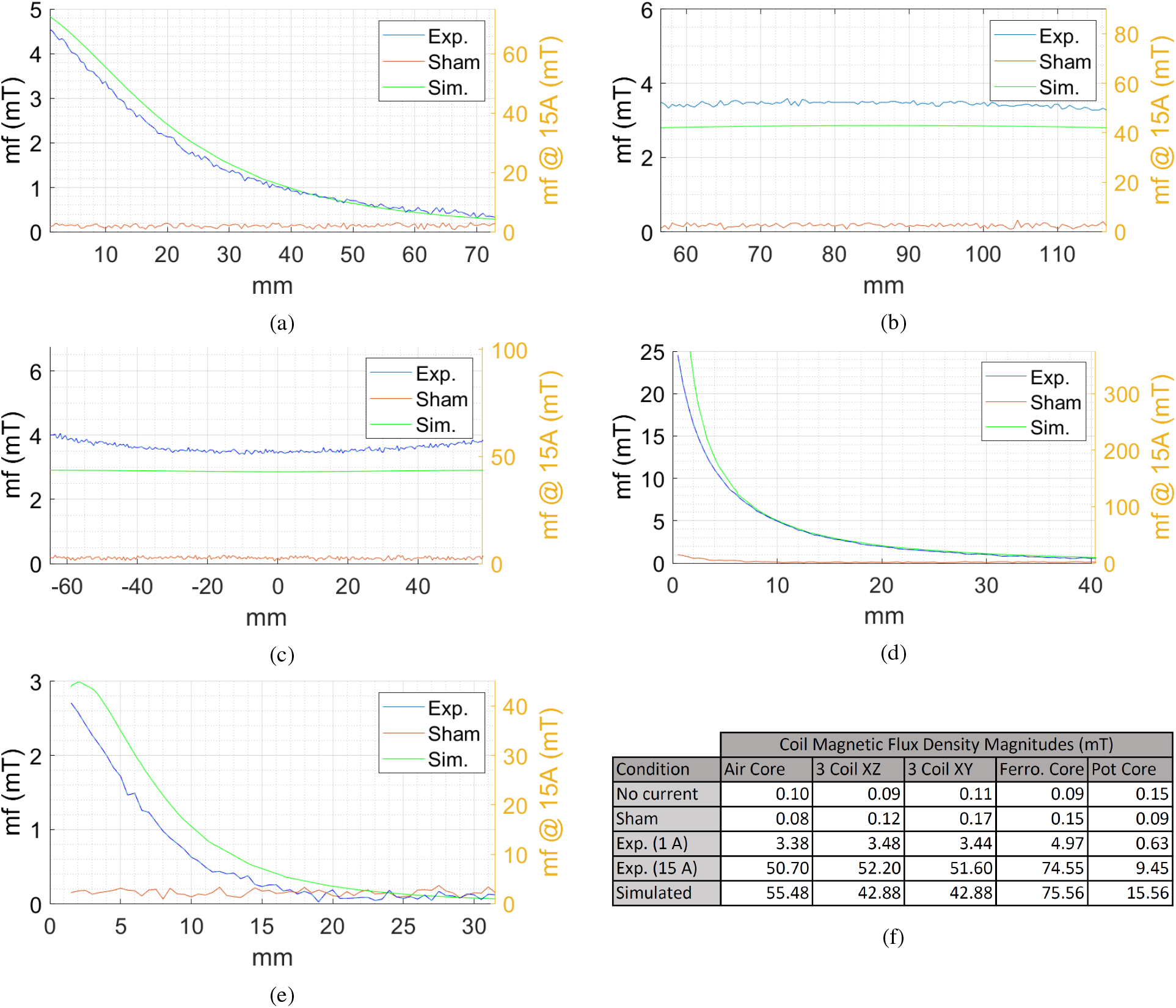
Line scans showing the experimental, sham, and simulated magnetic flux density measurements along the red line depicted in the corresponding distribution images of Figure 4 for the (a) air core coil, (b) three axis XZ, (c) three axis XY, (d) ferromagnetic core coil and (e) pot core coil scans. Experimental measurements were performed under a 1 A stimulation, simulated measurements were performed with 15 A, and the left and right axes of the graphs show the magnetic flux density for a 1 A and 15 A stimulation respectively. (f) Summary of the magnetic flux density magnitudes for the control, sham, experimental and simulated conditions. Measurements are from a distance 10 mm from the coil along the central axis for the air, ferromagnetic, and pot core coils and at the center for the three-coil system.

With regard to the uniformity of the stimulus delivered by the three-coil system, Figure 5b shows that the flux density along the three-coil system’s XZ axis drops on average only 2.99% in strength at the extrema of the line scan compared to the center. In the XY plane, the magnetic flux density increases on average 5.98% at ± 40 mm from the center, whereas at the extrema of the line scan it increases on average by 14.33% compared to the center.

It is worth noting that the ferromagnetic core does retain a low level of magnetization. However, the rapid decay in magnetic flux density means the magnetization has little effect at a distance of 10 mm from the tip.

Thermal imaging was also performed with a FLIR One thermal infrared camera. Safe operation of stimulation coils requires identification of maximum operating times for each coil geometry to stay below 75 °C. Such analysis is important to carefully design experiments that will allow the coils to stay within the defined temperature limits. Currents ranging from 1 to 15 Amperes at 1 Ampere intervals were applied to stimulate each coil while sampling coil temperature at 1 sample per second until the coil temperature reached 75 °C.

Results plotted in Figure 6 can be used to determine both the maximum excitation current and stimulation time based on the desired stimulus strength. For each of the four geometries, composite plots of maximum flux density magnitude and maximum operating time are shown for increasing stimulation current values ranging from 1 to 15 Amperes. The blue y-axis on the left shows the flux density magnitude in mT and the orange y-axis on the right shows the maximum operating times for the given stimulation current. Stimulation times are cutoff after one hour for the three-coil system and three minutes for the remaining geometries. An exponential best fit line for the maximum stimulation times is also shown in each graph of Figure 6.

**Figure 6:**
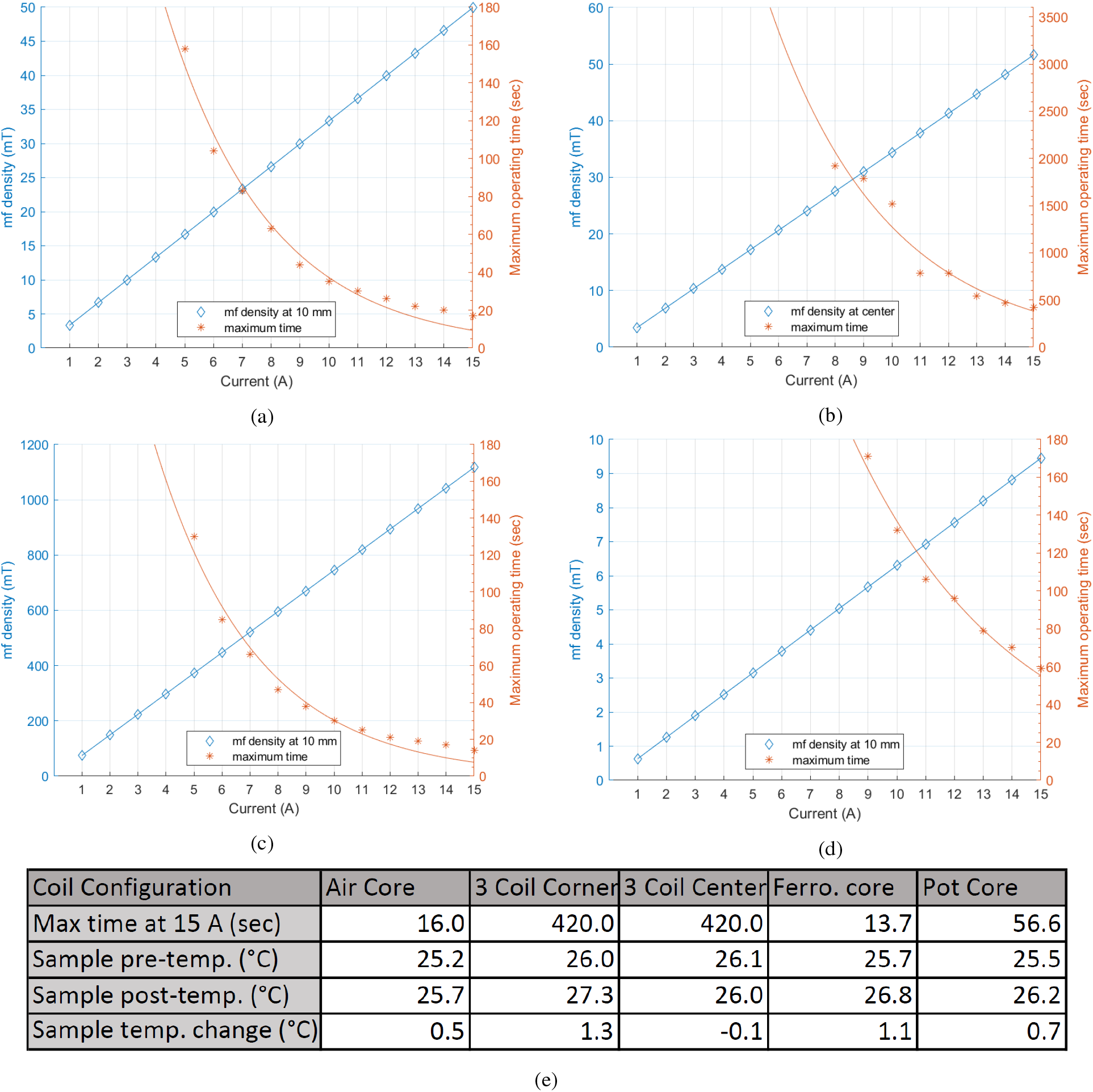
Magnetic flux density magnitudes, at 10 mm from the coil or the center for the three-coil system, shown on the left axis for a range of 1 to 15 Amperes for the (a) air core coil, (b) three-coil system, (c) ferromagnetic core coil, and (d) pot core coil. Also shown is the maximum operating time along the right axis for the same conditions. Operating times are shown only up to one hour for the three-coil system and three minutes for the remaining geometries. (e) Summary of the device operating times and sample temperatures for each coil design.

Temperatures were also measured at sample target locations for each coil configuration with a 15 Ampere stimulation current. Target locations for temperature measurements were the same locations as the measurements from Figure 5f. Additionally, temperature measurements were taken at the corner wells of a 96 well plate placed in the three-coil system. For the three-coil system, temperature was observed after application of a five minute stimulation. The air core, ferromagnetic core, and pot core coils were stimulated for the duration of their respective maximum operating times indicated in Figure 6.

With the air core coils, sample temperature was seen to rise by 0.5 °C, whereas the ferromagnetic core sample temperatures increased by 1.1 °C and the pot core sample temperatures increased by 0.7 °C throughout the stimulations. The temperature of the corner well samples in the three-coil system showed an increase of 1.3 °C after five minutes, while the temperature at the center remained essentially unchanged, decreasing by 0.1 °C.

## 6. Conclusion

This study presented a magnetogenetics stimulation platform that supports four electromagnet stimulation coil designs and a controller for selecting stimulation conditions. The various coil geometries were chosen so that at least one of the designs satisfied the needs of magnetogenetics experiments including microscopy, *in vivo* electrophysiology, freely moving behavioral experiments, and some fluorescence and luminescence imaging setups.

Regarding the use of ferromagnetic materials in stimulation coils, the added benefit of increased stimulation strength for an otherwise similar coil without a ferromagnetic core must be weighed against the necessity to account for the residual magnetization of the material. Negative effects can be mitigated by either demagnetizing the core between stimulations or placing the coil at a distance such that the residual field of the core does not interfere with the experiment design.

While sham conditions are required to ensure proper experimental controls, it is important to understand their limitations as they apply to a given experimental protocol and stimulation conditions. The sham conditions all demonstrated at least an order of magnitude reduction in magnetic flux density magnitudes, however, some residual magnetic flux is unavoidable. To properly incorporate sham conditions into an experiment, it would be best to know the minimum stimulation threshold necessary to produce a meaningful target response. With this knowledge, stimulations can be performed such that active conditions provide suprathreshold stimulations while sham conditions provide only subthreshold stimulation.

Accounting for the effects of temperature change is important when studying pathways with thermal sensitivity. Our designs showed minimal temperature increases (0.5-1.1 °C) at sample locations under maximum field strength conditions in all cases studied.

The versatility of various magnet designs presented allows for multiple choices of electromagnets based on the size constraints of the application. The analysis of the magnetic flux density distributions is important for selecting an appropriate electromagnet system to achieve the proper strength of the applied stimulus at the target location to successfully elicit a response. Additionally, our analysis of the sham stimulus strength is important in designing experiments with a negative control which can help eliminate the role of confounding variables on observed effects.

Studies presented here also provide a useful tool for selecting experimental design parameters for magnetogenetics experiments. For example, knowing that the three-coil system has a field distribution that varies less than 6% over a range of 40 mm from the central axis of the coils means that the sample placement should be restricted to this range in order to maintain a high degree of stimulation uniformity. In addition to limiting the effects of temperature on experimental observations, thermal analysis also allowed for determination of safe operating limits for the coils. Figure 6 can be used to find the operating limits, in terms of time and current, for a desired magnetic flux density with each coil geometry. Lastly, the use of a custom stimulation controller allows for easily configurable stimulation patterns, in either the sham or experimental modes, which improves repeatability.

## Acknowledgements

This work was supported by the National Institutes of Health: R01NS072171 (GP), R01NS098231 (GP and AAG) and R01NS104306 (AAG).

